# Humpback whales of the Persian Gulf

**DOI:** 10.1101/185033

**Authors:** Seyed Mohammad Hashem Dakhteh, Sharif Ranjbar, Mostafa Moazeni, Nazanin Mohsenian, Hossein Delshab, Hamed Moshiri, Seyed Mohammad B. Nabavi, Koen Van Waerebeek

## Abstract

The humpback whale has long been considered a rare straggler into the Persian Gulf, however new evidence contradicts this concept. We here critically review published and new records for *Megaptera novaeangliae* occurrence in the Gulf for the period 1883-2017. Of eight authenticated records (6 specimens, 2 live-sightings), seven are contemporary cases while one is a mid-Holocene specimen from UAE. An additional four are possible but unsubstantiated reports. Four regional, current, range states are confirmed, i.e. Iran, Iraq, Kuwait and Qatar. Four of the five newly reported cases are from Iran’s coastal waters. We conclude that the Persian Gulf is part of the habitual range of the Arabian Sea humpback whale population, and has been since at least the mid-Holocene. It is unknown whether frequent passage occurs through the Strait of Hormuz or whether whales are (semi)resident. The low abundance of this endangered population and frequent deleterious anthropogenic events, particularly ship strikes and net entanglements, are cause for major concern. In view of its historical and taxonomic relevance, the formal description of *Megaptera indica* Gervais, 1883, from Iraq, now thought to be a subspecies *M. novaeangliae indica*, is here translated from French.

## INTRODUCTION

An unique, because non-migrating, population of humpback whales *Megaptera novaeangliae* inhabits year-round the northern Arabian Sea, including the Gulf of Oman (Baldwin and Salm, 1994; Baldwin, 1998; Baldwin *et al.*, 1999; Mikhalev, 1997; Minton et al., 2011; Pomilla et al., 2014). The strong upwelling off Oman allows the whales to feed locally and forgo annual migrations (Papastavrou and Van Waerebeek, 1998). The Arabian Sea humpback whale forms a highly discrete stock that may have been reproductively isolated from other populations for some 70,000 years (Pomilla *et al.*, 2014).

A recent study of nuclear markers (Kershaw *et al.*, 2017) was consistent with the view of a discrete Arabian Sea breeding stock and found to be highly genetically differentiated (FST 0.034-0.161; P < 0.01 for all comparisons). The distributional boundaries of that population are poorly known, but extend to the Irani coast in the north (Braulik *et al.*, 2010; Owfi *et al.*, 2016), at least to Pakistan in the northeast (Van Beneden, 1887; Mikhalev, 1997), and to the Gulf of Aden (Mikhalev, 1997; Slijper *et al.*, 1964) in the Southwest. Based on a working document presented to the IWC Scientific Committee (Dakhteh *et al.*, 2017), we here critically review the long presumed rare occurrence of humpback whales in the Persian Gulf and suggest possible implications for management.

In the core distribution area, off the coast of the Sultanate of Oman, humpback whales seem to be concentrated off the Island of Masirah, Gulf of Masirah, Halaniyat Islands and Kuria Muria Bay in the Arabian Sea, considering that greatest numbers of records are from these areas (Baldwin *et al.*, 1999; Minton *et al.*, 2011). Humpback whales are sometimes seen near Fujairah, eastern coast of the United Arab Emirates (UAE) in the western Gulf of Oman (Baldwin, 1995), while the most northerly report on that coastline is of an individual at Khor [Khawr] Fakkan in 1973, some 80 km south of the Strait of Hormuz (M. Barwani, pers.comm. in Baldwin *et al.*, 1999). Although reported for the Persian Gulf by seamen (Slijper *et al.* 1964), till recently there was but a single authenticated record, namely Gervais (1883). Due to the lack of further evidence, the humpback whale has long been considered a rare visitor to the Persian Gulf (Baldwin *et al.*, 1999; Baldwin, 2003; Mikhalev, 1997; Robineau, 1998). However, a series of new, substantiated records is contradicting this view. Here we chronologically document and critically examine published humpback whale records, and add previously unreported records, for the Persian Gulf (Arabian Gulf). Potential but unsubstantiated reports are also discussed.

## SUBSTANTIATED RECORDS IN THE PERSIAN GULF

### 1. Al-Basra Bay, Iraq

The earliest record in the Persian Gulf is a specimen-supported 19^th^ century stranding of an adult humpback whale at the Al-Basra Bay (then named ’baie de Basora’) in Iraq, described as the type specimen of *Megaptera indica* by French taxonomist Paul François Gervais in 1883. Binomial nomenclature for many cetacean species, including the humpback whale, was highly unstable and proliferative at the end of the 19^th^ century with the introduction of many synonyms based on individual variation (e.g. Van Beneden, 1887; True, 1904). The *M. indica* nominal species has been universally re-assigned as a junior synonym of *Megaptera novaeangliae* (Borowski, 1781) (e.g. Hershkovitz, 1966; Tomilin, 1967; Robineau, 1989; Clapham and Mead, 1999). While the subspecific status of the isolated population of the Arabian humpback whale is currently under debate, the identity of the Al-Basra Bay animal as a humpback whale is indisputable thanks to the detailed osteological description including species-diagnostic features (Gervais, 1883) and cranial evidence re-examined by cetologist Daniel Robineau (1989, 1998). Unique among baleen whales, only in *M. novaeangliae* are the coracoid (processus coracoideus) and acromial processes of the scapula absent, or are expressed as rudimentary tubercles (True, 1904; Brinkmann, 1967; Tomilin, 1967). Gervais (1883) unequivocally described such a scapula in the Iraqi whale (see Appendix), as well as reported metacarpalia and phalanges in *M. indica* as even more elongated than in a specimen of *Megaptera Boops* (= *M. novaeangliae)* despite the latter measuring 2m longer than the *M. indica* holotype. The whale’s broad, uniformly black baleen are also consistent with humpback whale (see Appendix). Gervais (1888) published a second paper on the same specimen which is occasionally, equivocally, cited as the formal *M. indica* description.

While Gervais (1883) had access to both cranial and post-cranial bones (Appendix), Robineau (1989, 1998) found only the calvaria (JAC 1883-2255) at the Laboratoire d’Anatomie Comparée of the Muséum national d’Histoire Naturelle in Paris. He measured the calvaria as 3m in length with the tip of the rostrum missing, which agrees with the condylobasal length (CBL) of an adult humpback whale of some 12 m length, e.g. CBL= 314cm for a 12.20 m humpback whale *(in* True, 1904); CBL=311cm for a 11.96 m skeleton *(in* Tomilin, 1967).

Gervais (1883) considered the Iraq specimen a rare, extralimital record originating from the Indian Ocean, a view prevailing till recently (Baldwin et al., 1999). Considering its taxonomic and historical relevance and because it may be hardly accessible to many readers, we here present (Appendix) both the original text in French and an annotated, quasi-literal English translation.

### 2. Kuwait Inner Harbour, Kuwait

Mörzer-Bruyns (1971; p.182) reported that a humpback whale stayed one week in the Kuwait Inner Harbour in the western Persian Gulf, where it finally died after being hit by the propeller of a manoeuvring ship. Evidently pre-1971, there is no indication of date. The Dutch Captain Willem Frederik Jacob Mörzer-Bruyns, during his 40 years at sea, made a reputation as an experienced field observer of whales and dolphins and published several scientific papers on cetaceans. His ability to correctly recognise a live or freshly dead humpback whale, arguably the most identifiable of all baleen whales, should be undeniable and hence we consider this record to be valid.

### 3. Khour Mousa, Iran

One of us (S.M.B. Nabavi) registered a live-sighting of a single adult humpback whale around Khour Mousa (also spelled: Khur Moosa) in the western Gulf at N30°6.8492’, E49°10.299’, on 18 July 1984. After some 20 hrs the whale was seen to leave the area for deeper waters of the Gulf. The encounter is supported by two good photographs, showing the diagnostic low dorsal fin with a long base and a leading hump, and the flukes with an irregular serrated edge (Figure 1). When surfacing, the whale exposed three deep incisive injuries transversal across its dorsal fin. This pathomorphology is consistent with a sharp force trauma such as following collision with a large-vessel propeller. The injuries appeared unhealed and the survivability of this whale was unclear.

**Figure 1.**
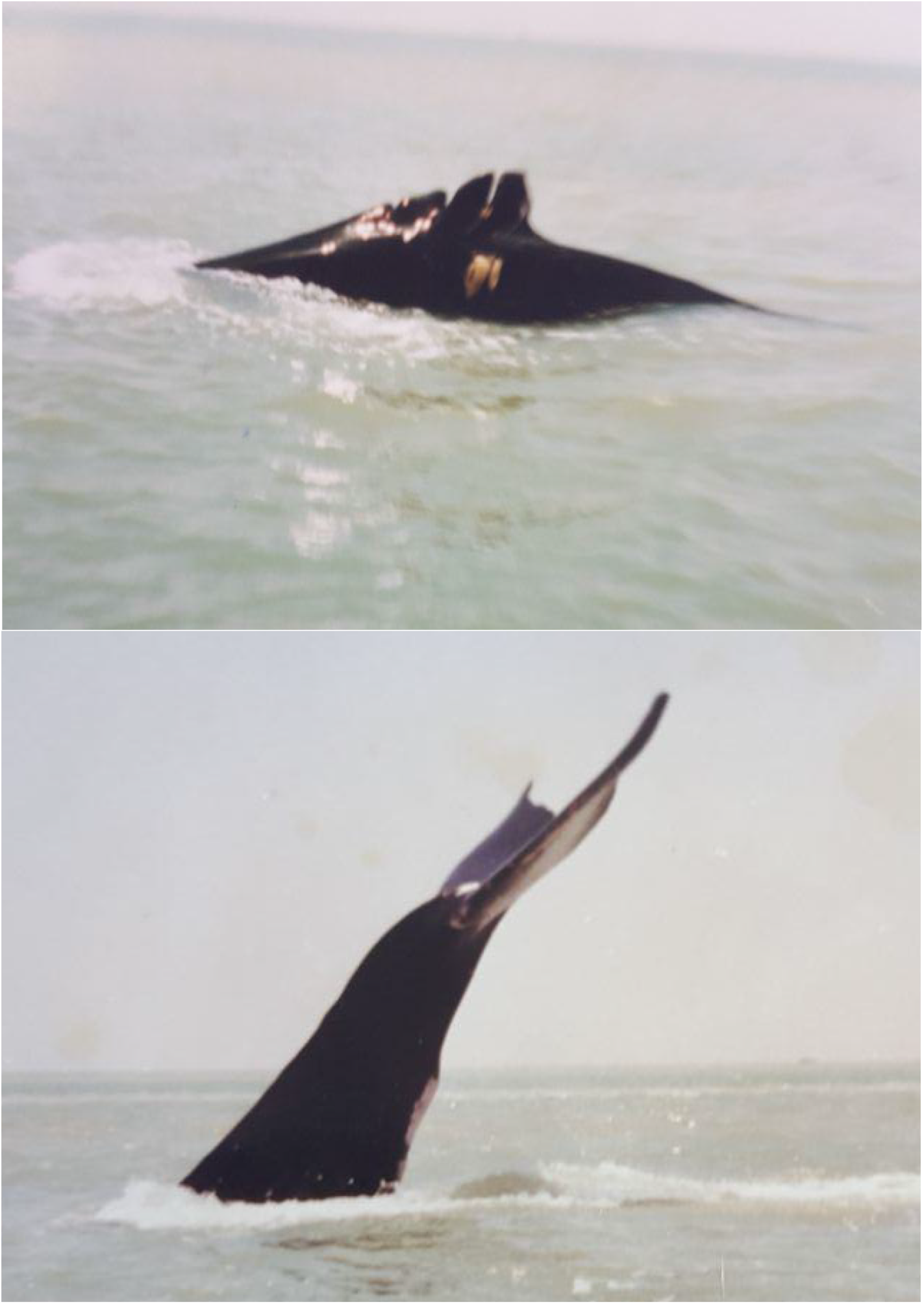
An adult humpback whale (record # 3) sighted near Khour Mousa, Iran, western Persian Gulf on 18 July 1984. Note deep injuries across the dorsal fin, consistent with sharp force trauma from large-ship propeller. ©Seyed M.B.Nabavi.

### 4. Bahrekan coast, Iran

The carcass of a juvenile humpback whale washed ashore on the Bahrekan coast in the shallow western Gulf, on 12 August 1996 and was examined by one of us (S.M.B. Nabavi). No exact position is available but the centroid of the Bahrekan coast is approximately at N30°05’,E49°42’. The body was relatively slender, grey in colour, with the dorsal fin set on a hump, the long right flipper measured ca. ¼ body length, the left flipper was severely damaged (Figure 2). The head had partly collapsed, presumably with loss of some cranial bones. The baleen plates were short. No samples were collected, but one low-resolution print photograph is available (Figure 2), which allowed a length estimate of 8.5-10 m deducted from the relative size of people standing beside the whale. In the field, Nabavi confidently identified it as a humpback whale, but the cause of death could not be established. The Bahrekan coast is an eutrophicated area subjected to organic pollution from wastes and heavy metals (e.g. Pb, Cu, Cd), hydrocarbons, urban wastewater pollution and biological impacts (i.e fisheries) (Shokat et al. 2010).

**Figure 2.**
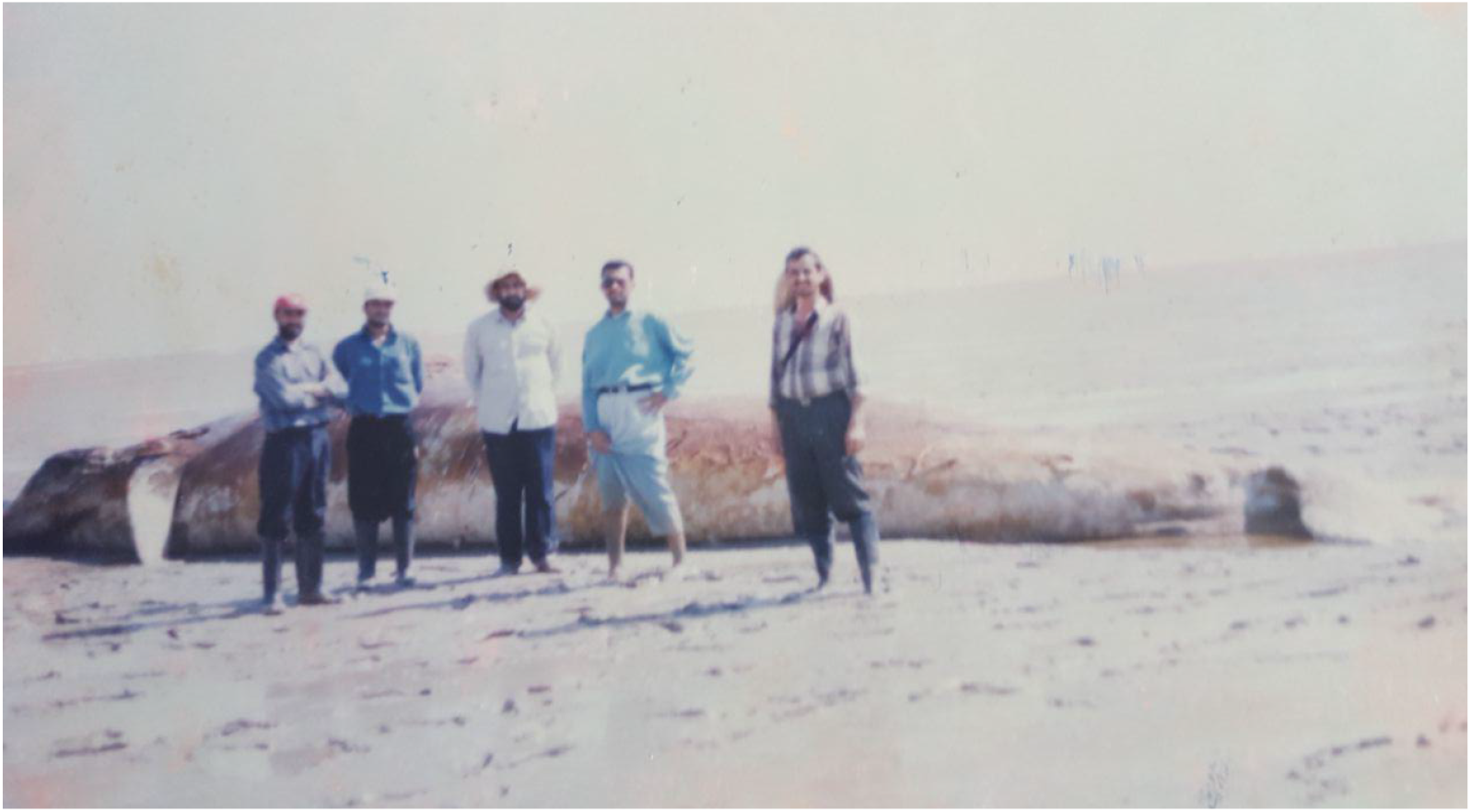
Juvenile humpback whale (record #4) stranded on Bahrekan coast, Iran, western Persian Gulf, on 12 August 1996. © Seyed M.B.Nabavi.

### 5. Doha, Qatar

Besides the Iraq record, Baldwin *et al.* (1999) in a comprehensive review mapped (their figure 10.2F) two unconfirmed (‘?’) humpback whale cases in the Persian Gulf, one from Qatar, the other from the United Arab Emirates (UAE). Later, Baldwin (2003) noted that the Iraq case is one of just two confirmed records from the Persian Gulf, leaving uncertainty about which was his second confirmed record. When queried by one of us (KVW), Robert Baldwin kindly contacted his original source, Dr. Tony Preen, who confirmed *(in litteris*, 17 Feb. 2017) a record in Qatar from a photo on display at the [National] Museum of Qatar which showed a dead humpback whale being lifted by a crane, presumably in the Doha port. While pre-1999, there is no date known for the photo. Considering that Dr. Preen is an experienced marine mammalogist we recognize this as a credible, photo-supported, humpback whale record.

### 6. Qeshm Island, Iran

Fishermen Mr. Abdolrahman Gurani and Mr. Yusof Poozideh, from Guran village, encountered a juvenile humpback whale entangled in their fishing net (Figure 3) in the channel between Khamir (Iran mainland) and Guran (Qeshm Island) on 6 July 2012. The site (N26°46.082’, E55°36.882’), locally known as Mosaageh, forms part of the Hara Mangrove Protected Area on the north coast of Qeshm Island, with estimated water depth some 8-9 m. The monofilament drift gillnet (length= 150m; depth= 5m; mesh size= 7.5 cm), set to target mainly silver pomfret *(Pampus argenteus)* of 1-6 kg each, typically is soaked for 8-9 days, checked regularly if not daily. As soon the owner-fisherman had been warned of the entanglement event by fellow fishermen, they proceeded to disentangle the whale to recuperate their net. The fishermen reported the size of the whale to be similar as their boat (6-7 m), corroborated by video which shows a smallish and thus juvenile humpback whale. The cellphone recorded video (.mp4) documented species-diagnostic characteristics, the head covered with the peculiar fleshy knobs (tubercles) and a dorsal fin set on a hump (Figure 3). Video voucher data are deposited at the Qeshm Environment Administration of the Qeshm Free Area, Qeshm City, Iran, and at the CEPEC library (Lima, Peru).

**Figure 3.**
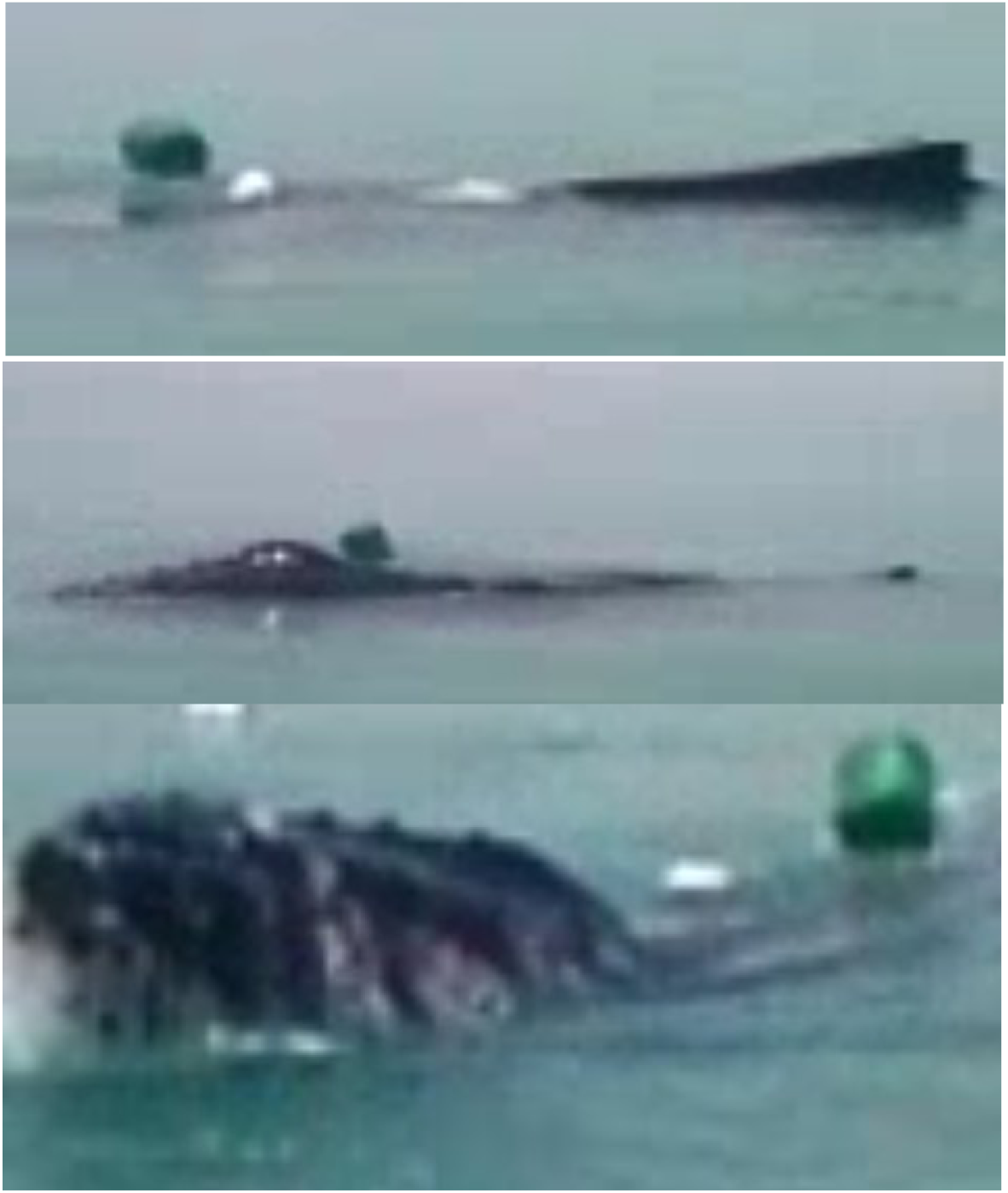
Three (non-successive) frames sampled from a cell-phone video, showing a gillnet-entangled juvenile humpback whale (record # 6) near Qeshm Island, Iran, in 2013. Although grainy, the frames unmistakably show the head with diagnostic fleshy knobs (tubercles) and a low, stubby dorsal fin with broad base. Video © Abdolrahman Gurani.

### 7. Akhtar, Iran

One of us (H. Delshab) first documented the relatively fresh, although bloating, carcass of a juvenile male humpback whale floating alongside the jetty of Akhtar Village (N27°41.722’, E52°11.6367’), Bushehr Province, on 19 April 2017 (Figure 4). The jetty was located on terrain owned by the South Pars Oil Company, near Kangan city. The unmeasured body length will be estimated once the skull is obtained. The cause of death is unknown but, considering juvenile age, is suspected to be anthropogenic, most likely shipping related. Photos showed no evidence of traumatic injuries ventrally, but no dorsal views were available. Upon discovery, due to strong wind and wave action the carcass could not be secured nor accessed for sampling, and over the next few days it drifted eastwards to wash ashore at N27°36.335’, E52°29.735’ close to Asalouyeh city, on 23 April 2017. Mostafa Moazeni, head of the Asalouyeh office of the Department of Environment (DoE), directed personnel to collect and bury the whale carcass for later retrieval of the skeleton. Other voucher data include photographs and a video deposited with the Plan for the Land Society, Tehran.

**Figure 4.**
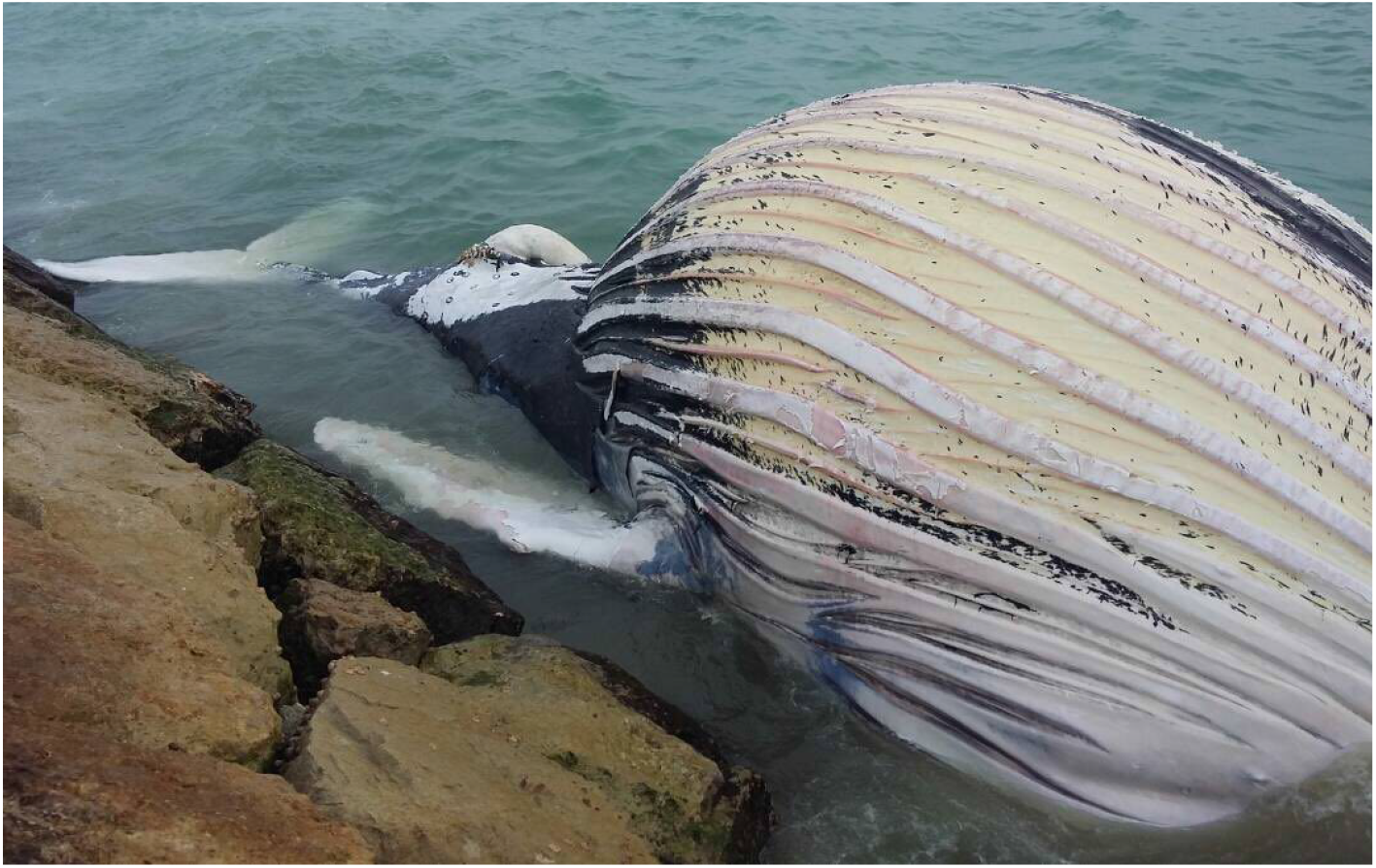
Freshly dead juvenile male humpback whale (record #4) floating at the jetty of Akhtar Village, Bushehr Province, Iran, on 23 April 2017.

**Figure 5.**
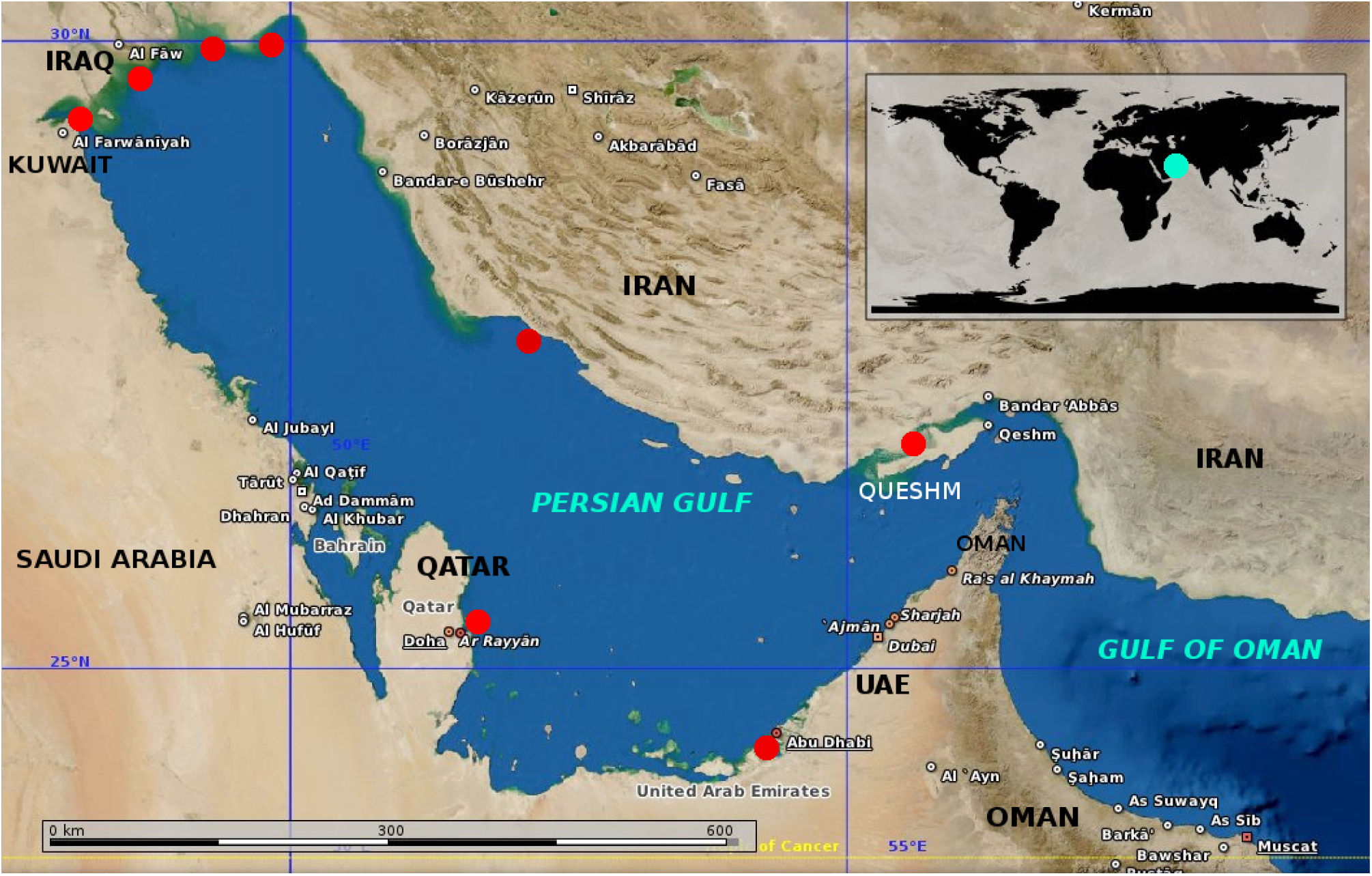
Distribution of eight substantiated records of humpback whales (red dots) in nearshore areas of the Persian Gulf [clockwise starting at Qeshm]: Qeshm Island,Iran; Musaffah, UAE; Doha, Qatar; Kuwait harbour, Kuwait; Al-Basra Bay, Iraq; Khour Moussa, Iran; Bahrekan Coast, Iran; Akhtar, Bushehr Province, Iran. No sightings are reported from offshore, international waters. Base map modified from Marble Virtual Globe.

### 8. Musaffah Industrial Channel, Abu Dhabi, UAE

Stewart *et al.* (2011) found whale remains (a left and right mandible, scapula, humerus and fragmentary radius and ulna as well as parts of the cranium and rostrum) belonging to a ’probable humpback whale (*Megaptera* cf. *novaeangliae)’* in the well-described sabkha sequence exposed in the Musaffah Industrial Channel, Abu Dhabi, UAE. More precisely, the whale remains were found in a series of sediments representing a range of lagoonal facies. The sediments surrounding the whale bones were age-dated (14C) at approximately 5200 yrs BP (Holocene) and are therefore interpreted to correspond to the previously documented late Flandrian sea-level peak, preceding a fall in sea-level which culminated in the supratidal sabkha overprint of the carbonates (Stewart *et al.*, 2011). *Megaptera novaeangliae* has existed since at least the latest Middle Pleistocene (Nagasawa and Mitani, 2004), in the western North Pacific (Japan), and the Arabian Sea humpback whale population is thought to have been isolated from others since 70,000 yrs (Pomilla et al., 2014).

The fan-shaped scapula without coracoid and acromion processes (see figure 6, *in* Stewart *et al.* 2011), the absent or reduced olecranon of the ulna, and the relatively more laterally directed zygomatic processes of the squamosal compared to *Balaenoptera* (Tomilin, 1967; Winn and Reichley, 1985; Deméré *et al.*, 2005) demonstrate that there is no doubt about its taxonomic identity as humpback whale.

## POSSIBLE BUT UNSUBSTANTIATED REPORTS

1. An unconfirmed report relates to a vertebra and a rib in the Iraqi Natural History Museum found about 1954. There is an old report that a Turkish gunboat killed this whale about a century ago in the Shatt-al-Arab’ (river dividing Iraq and Iran) (Al-Robaae, 1974; De Silva, 1987). The original source was R. Hatt, 1959 but was not seen. We could not verify whether these bones or any associated information still exist. In view of the positive Al-Basra Bay (#1) and Kuwait harbour (#2) records, also from the western Gulf, a humpback whale in the Shatt-al-Arab is plausible.

2. Slijper et al. (1964, their Chart 5) mapped three sightings of humpback whales in the Persian Gulf, one off the UAE in May and two at the extreme western end of the Gulf, in the period 1954-1956. However these observations were made by seamen, not biologists, and the lack of voucher material does not allow us to verify identifications. Slijper et al. (1964) believed they were reliable. It is likely that the Baldwin et al. (1999) ’possible record’ for the UAE was based on this information.

## DISCUSSION

After reviewing published and unpublished evidence we authenticated eight records of humpback whales in the Persian Gulf, seven contemporary cases and one mid-Holocene specimen. Four regional range states are confirmed, i.e. Iran (n=4), Iraq (n=1), Kuwait (n=1) and Qatar (n=1). No documented records exist from coastal waters of Saudi Arabia or Bahrain (Baldwin *et al.* 1999; Baldwin, 2003). The UAE yielded the subfossil record but although its central coast has been flagged as a potential location of recent occurrence (Slijper et al., 1964; Gallagher, 1991; Baldwin *et al.*, 1999), no hard evidence could be located (R. Baldwin, *in litteris* to KVW). Also, we found no published sightings for international waters of the Persian Gulf.

Four of five new records originated from Iran. Humpback whales are commonly cited in check-lists of the mammals of Iran (Firoz, 1976; Harrington, 1977; Etemad, 1984; Humphrey and Kharom, 1995; Ziaie, 1996; Firouz, 2005). Owfi *et al.* (2015) suggest that ‘these records appear to be based on *Balaenoptera* sp. skeletons that have been mis-identified as humpbacks’. However, until specimens are properly identified to species, unrecognised cases of *M. novaeangliae* may appear. Three reports for Iran’s Gulf of Oman coast (Braulik *et al.*, 2010; Owfi *et al.*, 2015) originate from the Sistan/ Baluchistan Province. These include a mother-calf pair sighted near Chabahar in September 2004, a stranding at Pozm (50 km W. of Chabahar) in October 2004 and a third stranding at an unspecified location in December 2003. Unfortunately no details or verifiable voucher materials were presented, customary for new range state records. Therefore the cases reported here actually represent the first fully documented records of *M. novaeangliae* in Iranian waters.

The Arabian Sea humpback whale is the only known population in the region. A number of records for the NW Gulf of Oman near the entrance of the Strait of Hormuz (Baldwin et al., 1999; Minton *et al.*, 2011; Pomilla *et al.*, 2014) allow us to reasonably assume a single-stock continuous distribution into the Gulf. Tissue samples for molecular genetics should ascertain this. The mid-Holocene specimen from UAE (Stewart *et al.*, 2011) indicates that humpback whale presence in the Persian Gulf is not the result of any recent ecological or climatic changes, but that the Gulf has long been part of the habitual range of the Arabian Sea population.

Hopefully, future more systematic marine mammal surveying will also collect environmental data which should shed light on the factors that make the Persian Gulf a suitable long-term habitat, if not permanent residence, for humpback whales. This shallow sea may offer favorable feeding or reproductive conditions, or both. In the Gulf of Oman, humpback whales sometimes enter very shallow water to feed on schools of sardines, anchovies, chub mackerel, scad and similar small fishes (Baldwin and Salm, 1994; Baldwin, 2003). Whales in the Persian Gulf may be both piscivore or prey on euphausids. Surprisingly perhaps, but high densities of the euphausid *Pseudeuphausia latifrons* (200-299 and > 300 individuals/100 m3) are reported from the northern Persian Gulf off Iran (Weigmann, 1970a,b). In six of the seven reported cases in the Gulf, the whales were encountered in shallow, nearshore waters. The net-entangled juvenile near Qeshm Island (#6) is thought to have been feeding. Predators may also influence distribution. The killer whale (*Orcinus orca)*, a known predator of humpback whales (Clapham and Mead, 1999), has been mentioned for the Persian Gulf (Baldwin, 1995, 2003; Ada Natoli, *in litt.* to KVW, 23 August 2017), however no specific records have been published (Owfi et al., 2016) and they are thought to rarely enter the Gulf. If so, this might offer an additional incentive for humpback whales to reside in the Gulf.

The deleterious anthropogenic effects on humpback whales in the Persian Gulf are of major concern. Of the seven confirmed contemporary records (i.e. excluding mid-Holocene specimen), only two whales were seen alive, one of which was net-entangled (#6) and the second (#3) was severely injured by a propeller collision. Among the five dead ones, at least three were juveniles that suffered an untimely death. One was a confirmed collision case (#2), and three were probable collision victims (#1, 5, 7) as they were suspiciously found inside a port or in the general vicinity of a portuary area. In addition, albeit an unconfirmed record, a whale was killed by a gunboat in Iraq. Other possible threats may include chemical contamination and oil spills (e.g. Robineau and Fiquet, 1994; Preen, 2004) and infectious diseases (Van Bressem *et al*., 2014).

One might argue that four of the whales may have died considerable distance from location of reporting. For instance, theoretically, the Qatar specimen (#2) might have been struck and killed by a large ship, and transported, wrapped on the ship’s bulbous bow, many tens or even hundreds of km from Doha. Except that, while *M. novaeangliae* globally is the second-most commonly killed whale species by ship collisions, unlike other balaenopterids the species remains rarely stuck on the bow of vessels (Van Waerebeek *et al.*, 2007), probably because of its axially asymmetric body shape. Only very few vessels (only military) had a bulbous bow in the late 19^th^ century, and we suggest that the Iraq whale (#1) probably died near Bassora Bay.

Morphological (Gervais, 1883; Appendix 1), genetic (Minton *et al.*, 2011; Pomilla *et al.*, 2014), behavioural (Whitehead, 1985) and distributional (for non-migratory) (Mikhalev, 1997; Papastavrou and Van Waerebeek, 1998) lines of evidence concord that the Arabian Sea humpback whale population has been reproductively isolated from other populations sufficiently long to differentiate. Enhanced genetic drift within a small population may accelerate the speciation process, and we agree with Pomilla *et al*. (2014), that the Arabian Sea population deserves subspecific status, *Megaptera novaeangliae indica.* A comparative-morphological study with new specimens should re-examine earlier findings. Gervais (1883) may have emphasized disproportionate importance to variation seen in the sternum and tympanic bulla. Sternal morphology is highly variable intraspecifically in Mysticeti, and its variability is so great that it can hardly be used as a criterion for the separation of species (Omura, 1975; Klima, 1978). Sterna in humpback whales may be triangular, heart-shaped, trilobate or U-shaped (Klima, 1978).

The world’s so-called ’most isolated humpback whale population’ combined with very low, declining, population abundance (82 ind., 95%CI 60-111 from capture-recapture; 90-142 ind. from genetic data) (Pomilla *et al.*, 2014) raise extreme concern for this population’s continued survival. Soviet whaling data suggested still at least 400 individuals 50 years ago (Pomilla *et al.*, 2014). Dedicated marine mammal research throughout the Persian Gulf and the northern Arabian Sea should be augmented, and besides the recording of biological data also information on human-caused mortality and morbidity, which means predominantly fisheries interactions and ship strikes. Finally, epidemiology of emerging infectious diseases also deserve priority attention (e.g. Van Bressem *et al.*, 2014), as the strong isolation may mean that members of this population may be immunologically naïve and therefore highly susceptible to certain lethal epizootics.

## ACKNOWLEDGEMENTS

We are very grateful to Mr. Abdolrahman Gurani and Mr. Yusof Poozideh of Guran village for voluntarily informing the Qeshm Environment Administration about whale entanglement; to Drs. Tony Preen and Robert Baldwin for kindly responding to various queries in our attempt to elucidate Qatar and UAE reports. We warmly thank Mrs. Laleh Daraie (SGP coordinator in Iran), Mr. Houman Jokar and Mrs. Sepideh Kashani for contributing to various activities by the Qeshm Environment Administration, as well as Mr. Amin Tollab (Marine Environment Bureau of the Department of Environment of Bushehr Province) and Mr. Ahmad Azaryar (Head of Kangan office, Department of Environment, Bushehr Province). Many thanks are due also to Mrs. Dr. Farshchi (Deputy of Marine Environment of DOE) and Dr. Mirshekar (General Director of Marine Ecosystems of DOE) for their most helpful collaboration.

The paper has benefitted from much appreciated exchanges of thoughts on cetaceans of the study region with Drs. Ada Natoli, Robert Baldwin and Gianna Minton. Drs. Anand Manickam, Simon Wilson and Laith Jawad are thanked for literature support. The Small Grants Programme (SGP) of UNDP/GEF in Iran and the Persian Wildlife Heritage Foundation co-supported work by the Qeshm Environment Administration. Plan for the Land Society thanks the ASHW network for advice and the fishermen and residents of Akhtar village for their help.

## APPENDIX On a new species of the genus *Megaptera*, originating from the Bay of Basora [Al-Basra] (Persian Gulf) by H.-P. Gervais (1883)

» Translated and annotated [in square brackets] by one of us (KVW.

» The genus *Megaptera*, as established by the authors of the *Ostéographie des Cétacés* [i.e. Van Beneden & Gervais, 1880] comprehends four distinct species; the first two, the *Megaptera Boops* [(Fabricius, 1780)] and *Megaptera Lalandii* [(Fisher, 1829)] are established with certainty; the two others, *Megaptera Novae-Zelandiae* [Gray, 1864] and *Megaptera Kuzira* [Gray, 1850] are described only provisionally as their characteristics are still insufficiently clear.

» Although Mr. [Pierre-Joseph] Van Beneden, in a recent work, has come back from the idea that he formulated 20 years ago that there exists but a single cosmopolitan species of *Megaptera*, namely *Megaptera Boops*, we think that we can demonstrate from a study that we undertook, through the comparison of new material accumulated at the anatomical collections of the Paris Museum, that the law of species distribution established for the [families] Balaenidae and Balaenopteridae, has to apply also to *Megaptera* and that the number of species of this group have to be recognised as three. These are the *Megaptera Boops*, inhabiting the Northern Hemisphere; the *Megaptera Lalandii* which frequents the southern part of the Atlantic Ocean, and the *Megaptera* of the Persian Gulf, which is the subject of the present note, a species which is supposed to inhabit the Indian Ocean and to which we propose to give the name *Megaptera indica*, since the individual obtained for the collections of the Museum would only have penetrated the Persian Gulf accidentally, from where it was shipped to us.

» The size of the humpback whale of the Persian Gulf, which had attained adult age, differs hardly from the size of the skeleton of the equally adult *Megaptera Boops* with which we compared it. The external body shape should however have been more slender and its head more rounded.

» The general shape of the bony head shows, in its dorsal contours, a much more masked curvature: the rostrum is more obtuse, the lower part of the maxillary is more arched. The occipital region of the skull is less concave than in the *Megaptera* from the north, the longitudinal crest occupying the middle of the external face of the occipital is more pronounced. The lateral occipital protrusions are more marked while the condylar region is less prominent; the occipital opening [foramen magnum] is located less high and, consequently, projects more to the rear.

» The os temporale differs especially in its zygomatic part, which is shorter, more massive, more arched at its top and is more outwards projecting. The frontal bones demonstrate also, in their shape, fairly large differences; their orbital extensions are more massive, in a less oblique way from inside to outside and from the rear to anterior. The optical groove is largely open over its entire tract.

» The lower part of the skull, although somewhat damaged, has nonetheless permitted us to note that the os palatinum, which is highly characteristic with respect to the distinction between cetacean species, differs in shape, shows more considerable thickness and their large articulation with the maxillaries [’maxillaires supérieurs’] in the *Megaptera* from the Persian Gulf. The pterygoid bones are also very heavy, and their posterior apophysis, much shorter and bulkier than in *Megaptera Boops*, is strongly recurved towards the rear and outwards.

» The maxillaries’ external borders are less straight than in the northern species. The rostrum shows a fairly marked narrowing somewhat anterior to the base of the orbital apophyses, then it widens in the middle region before progressively narrowing towards its anterior end. All the parts show, rather pronounced, different characteristics.

» The os jugale and the os lacrimale also show a particular configuration in our animal.

» The vertebrae overall are distinguished by the thickness of their bodies [corpus vertebrae], which is more pronounced in the first few cervicals of the *Megaptera* of the Persian Gulf than the corresponding bodies in the *Megaptera* of Lapome [= *M. Boops]*, which however was of greater length. The transverse and spinous processes in the former species, compared to the latter species, differ in shape and direction; they are generally shorter, broader and thicker. The transverse processes of the dorsal region, especially those that occupy the middle part of that region, are higher than can be seen in any of the skeletons described till now and bring our *Megaptera* closer to the right whales, more than any other species of the group.

» The first and the second cervical vertebrae deserve to be especially mentioned: the atlas is distinguished from the one in *Megaptera Boops* by the curvature of its upper neural arch, the thickness of its [vertebral] lamina of which the posterior border is grooved by two deep articular cavities [facets] into which the two articular apophyses fit from the anterior rim of the neural arch of the axis; the upper transversal processes are shorter and more massive.

» The lower transversal processes are well-developed, which is not the case in *Megaptera Boops;* the process at the right is ankylosed with the upper transversal process and forms an apophyseal mass, apparently unique, at which base one can find a large vertebral canal [foramen].

» The second cervical vertebra or axis differs as much from that of the northern species as the two atlas vertebrae are different between them. The third cervical vertebra shows two pairs of highly developed transversal processes, namely upper and lower.

» The thoracic limb [flipper] is longer than the one in *Megaptera Boops*, with which we compare it, even while considering that the body size of the first one is 2 meter shorter than the second one. The scapula does not bear an acromion [diagnostic for *Megaptera];* the coracoid process is represented by a small bone protrusion [diagnostic for *Megaptera]* and the overall shape of this bone differs measurably between the two species. All metacarpal bones are larger, longer and thicker than in the northern species; they contribute jointly with the plalanges which are also longer, larger and more flattened to give to the flipper of this animal its larger dimensions.

» The ribs are less long and more rounded than in the other two species.

» The *sternum* in our *Megaptera* from the Persian Gulf differs completely in shape from all the Mysticete species described so far. This bone is relatively very small, although we are dealing with an adult specimen. Its shape is like some sort of tail flukes of which the anterior face is concave in vertical sense and convex transversally. The lateral extensions, articulating with the first pair of ribs, are barely noticeable. All the edges are rounded, especially the anterior one which is thick and curved forward; the lower border ends in a pronounced triangular point.

» The tympanic bulla of the *Megaptera* of the Persian Gulf has a characteristic shape; and is remarkable by its small dimensions.

» The baleen plates are large, thick and coloured uniformly black.

## Sur une nouvelle espèce du genre Mégaptère, provenant de la baie de Basora (golfe Persique); par M. H.-P. Gervais

» Le genre Mégaptère, tel qu’il a été établi par les auteurs de *l’Ostéographie des Cétacés*, comprend quatre espèces distinctes; les deux premières, la *Megaptera Boops* et la *Megaptera Lalandii*, y sont établies d’une façon certaine; les deux autres, la *Megaptera Novae-Zelandiae* et la *Megaptera Kuzira*, n’y sont inscrites que d’une façon provisoire, leurs caractères étant insuffisamment connus. » Bien que M. van Beneden, dans un récent Ouvrage, soit revenu à l’idée qu’il avait émise il y a plus de vingt ans, qu’il n’existerait qu’une seule espèce de Mégaptères cosmopolite, la *Megaptera Boops*, nous croyons pouvoir, par la comparaison de nouveaux matériaux rassemblés dans les collections anatomiques du Muséum de Paris, démontrer, dans un travail que nous avons entrepris, que la loi de répartition des espèces établie pour les Balaenidés et les Balaenoptères doit s’appliquer aussi aux Mégaptères et que le nombre des espèces de ce groupe doit être porté à trois, qui sont: la *Megaptera Boops*, habitant ľhémisphère boréal; la *Megaptera Lalandii*, fréquentant la partie sud de l’océan Atlantique, et la *Megaptera* du golfe Persique, qui fait le sujet de la présente Note, espèce qui habiterait l’océan Indien et à laquelle nous proposons de donner le nom de *Megaptera indica*, car ce n’est qu’accidentellement que l’individu acquis pour les collections du Muséum aurait pénétré dans le golfe Persique d’où il nous a été expédié.

» La taille de la Mégaptère du golfe Persique, qui est arrivée à l’âge adulte, diffère à peine par le squelette de la *Megaptera Boops*, également adulte, à laquelle nous avons pu lui comparer. Les formes extérieures de son corps devaient être pourtant plus élancées et la tète plus globuleuse.

» La forme générale de la tête osseuse accuse, dans ses contours supérieurs, une courbure beaucoup plus masquée: le rostre est plus obtus, le maxillaire inférieur plus arqué. La région postérieure du crâne est moins concave que chez la Mégaptère du Nord, la crête longitudinale occupant le milieu de la face externe de l’occipital est plus accentuée, les saillies des occipitaux latéraux plus marquées et la région condylienne moins proéminente; le trou occipital est situé moins haut et regarde, par conséquent, plus en arrière.

» L’os temporal diffère surtout dans sa portion zygomatique, qui est plus courte, plus massive, plus arquée à son sommet et dirigée plus en dehors. Les os frontaux accusent aussi, dans leur forme, des différences assez grandes; leurs prolongements orbitaires sont plus massifs, à direction moins oblique de dedans en dehors et d’arrière en avant. La gouttière optique est largement ouverte dans toute l’étendue de son trajet.

» La région inférieure du crâne, quoique un peu mutilée, nous a permis pourtant de remarquer que les os palatins, qui donnent de si bons caractères, au point de vue de la distinction des espèces, chez les Cétacés, diffèrent par leur forme, leur épaisseur plus considérable et leur large articulation avec le maxillaire supérieur chez la Mégaptère du golfe Persique. Les ptérygoïdiens sont aussi très épais, et leur apophyse postérieure, beaucoup plus courte et plus forte que chez la *Megaptera Boops*, est très recourbée en arrière et en dehors.

» Les maxillaires supérieurs ont leurs bords externes moins droits que chez l’espèce du Nord. Le rostre subit un rétrécissement assez marqué un peu en avant de la base des apophyses orbitaires, puis il s’élargit dans sa région moyenne pour diminuer ensuite progressivement vers son extrémité antérieure. Toutes leurs parties présentent des caractères différents assez marqués.

» L’os jugal et l’os lacrymal ont aussi une configuration particulière chez notre animal.

» Les vertèbres se distinguent d’une façon générale par l’épaisseur de leur corps, qui est plus grande dans les premières cervicales chez la Mégaptère du golfe Persique que celles qui leur correspondent dans la Mégaptère de Lapome, qui était pourtant supérieure quant à la taille. Les apophyses transverses et épineuses de la première de ces espèces, comparées à celles de l’autre, diffèrent comme forme et comme direction; elles sont généralement plus courtes, plus larges et plus épaisses. Les apophyses transverses de la région dorsale, surtout celles qui occupent le milieu de cette région, sont plus relevées que cela ne se voit chez aucun des squelettes décrits jusqu’ici et rapprochent notre Mégaptère, plus qu’aucune autre espèce du groupe, des vraies Baleines.

» Les deux premières vertèbres cervicales méritent une mention spéciale: l’atlas se distingue de celui de la *Megaptera Boops* par la courbure de son arc supérieur, l’épaisseur de ses lames dont le bord postérieur est creusé de deux cavités articulaires profondes dans lesquelles pénètrent deux apophyses articulaires venant du bord antérieur de l’arc neural de l’axis; ses apophyses transverses supérieures sont plus courtes et plus massives.

» Les apophyses transverses inférieures se sont développées, ce qui n’a pas lieu chez la *Megaptera Boops;* celle du côté droit, soudée avec l’apophyse transverse supérieure, forme une masse apophysaire en apparence unique, à la base de laquelle se trouve un large canal vertébral.

» La deuxième vertèbre cervicale ou axis diffère autant de celle de l’espèce du Nord que les deux vertèbres atlas diffèrent entre elles. La troisième vertèbre cervicale porte deux apophyses transverses supérieures et inférieures très développées.

» Le membre thoracique est plus long que chez la *Megaptera Boops* à laquelle nous la comparons, bien que la taille du premier de ces sujets soit inférieure de près de 2 m à celle du second. L’omoplate est dépourvue d’acromion; l’apophyse coracoïde est représentée par une petite saillie osseuse et la forme générale de cet os diffère sensiblement chez les deux espèces. Tous les métacarpiens sont plus larges, plus longs et plus épais que dans l’espèce du Nord; ils contribuent, avec les phalanges qui sont aussi plus longues, plus larges et plus aplaties, à donner à la nageoire de cet animal de plus grandes proportions.

» Les côtes sont moins larges et plus arrondies que dans les deux autres espèces.

» Le sternum, chez notre Mégaptère du golfe Persique, diffère complètement par sa forme de celui de toutes les espèces de Mysticètes décrites jusqu’ici. Cet os est relativement très petit, quoique nous ayons affaire à un sujet adulte. Sa forme est celle d’une sorte de battoir dont la face antérieure est concave dans le sens de la hauteur, convexe transversalement. Les prolongements latéraux, s’articulant avec la première paire de côtes, sont à peine sensibles. Tous les bords, surtout le bord antérieur, qui est épais et recourbé en avant, sont arrondis; le bord inférieur se termine par une forte pointe triangulaire.

» La caisse tympanique présente chez la Mégaptère du golfe Persique une forme caractéristique; elle se fait remarquer par ses faibles dimensions.

» Les fanons sont larges, épais et de couleur uniformément noire. »

(31 décembre 1883).

